# Complete ammonia-oxidizing *Nitrospira* and ammonia-oxidizing archaea as biological supplements for freshwater aquaria

**DOI:** 10.64898/2026.10.08.757816

**Authors:** Alexander K. Umbach, Owen Ikert, Benoit Choquet, Josh D. Neufeld

**Author notes:** Corresponding author: Department of Biology, University of Waterloo, 200 University Avenue West, Waterloo, Ontario, N2L 3G1, Canada.

## Abstract

Biological filtration in newly established aquaria depends on the establishment of nitrifying microorganisms that prevent accumulation of toxic ammonia and nitrite. Many aquarists use commercial nitrifying supplements to accelerate biofilter establishment and reduce ammonia and nitrite accumulation. However, the microbial composition and *in situ* efficacy of these products remain poorly characterized. We evaluated the ability of enrichment cultures containing comammox *Nitrospira* (CX) and the ammonia-oxidizing archaeon *Ca.* Nitrosotenuis aquarius (AQ) to serve as aquarium nitrifying supplements, tested alongside a conventional supplement (CS) containing ammonia-oxidizing bacteria. Inoculation of new aquaria containing zebrafish (*Danio rerio*) with nitrifying supplements resulted in distinct nitrifier profiles within biofilters of each treatment group corresponding with significant reductions in peak ammonia and nitrite concentrations, which were most evident when applied at the maximum tested supplement dose. Treating aquaria with CX, AQ, or CS maintained ammonia and nitrite at low concentrations, and CX effectively prevented ammonia and nitrite accumulation through all stages of aquarium operation and biofilter establishment. After removing all fish from the aquaria, ammonia “spike-in” experiments demonstrated biofilters treated with CX and AQ conferred long-term protection against nitrite accumulation following acute ammonia loading events. Whereas 16S rRNA gene sequences associated with CS were rarely detected within biofilter profiles, those associated with CX and AQ established early and persisted temporally. Overall, results demonstrate how nitrifiers might be differentially adapted to home aquarium conditions and the potential of AOA and comammox *Nitrospira* to improve biofilter establishment and maintaining safe water quality in newly established freshwater aquaria.

**Importance:** Establishing nitrifying microbial populations in freshwater aquarium biofilters is required for maintaining water quality and preventing ammonia and nitrite toxicity. Although new aquaria may be seeded by commercial nitrifying supplements, the microbial composition and *in situ* efficacy of these cultures remain poorly characterized. This study tests the ability of aquarium-derived enrichment cultures of comammox *Nitrospira* and the ammonia-oxidizing archaeon *Candidatus* Nitrosotenuis aquarius for use as nitrifying supplements. Results demonstrate that both cultures augment nitrification within newly established freshwater aquaria, especially at increased dosages. Compared to an existing supplement containing ammonia-oxidizing bacteria, the comammox-containing enrichment maintained ammonia and nitrite oxidation activity within filters to prevent ammonia and nitrite accumulation. These results connect microbial community establishment with a practical bioaugmentation application and demonstrate the suitability of supplement microorganisms to establish and persist within biofilter biofilm and reveal that inoculum size is an important consideration for validating nitrifying cultures for supplement application.

## Introduction

When establishing a new freshwater aquarium, accumulation of harmful ammonia (NH_3_) and nitrite (NO_2-_) is a concern because these compounds become toxic to aquatic organisms under a range of concentrations, some as low as 0.1 mg/L-N (1–3). In established aquarium biofilters, a nitrifying consortium of ammonia-oxidizing archaea (AOA), ammonia-oxidizing bacteria (AOB), nitrite-oxidizing bacteria (NOB), and comammox *Nitrospira* is suspected to oxidize ammonia and nitrite (4–6). However, new aquarium biofilters are established with new filter media that is largely absent of microbial communities and their associated nitrifying guild members. Fish added to a new aquarium are thus susceptible to “new tank syndrome” (7, 8), a term describing the accumulation of ammonia in the absence of sufficient nitrifying activity, resulting in fish stress, illness, or death. A common strategy among aquarists for avoiding new tank syndrome is to “cycle” a new aquarium, a process commonly detailed in online forums and pet websites (9–13). Briefly, cycling an aquarium relies on passive inoculation and enrichment of nitrifiers within the biofilter in the absence of fish, or with relatively few fish added. Ammonia and nitrite concentrations are monitored over the course of several weeks until they are below detectable limits and fish are then added with the assumption that sufficiently abundant nitrifying populations have been established.

An alternative approach for establishing a new aquarium biofilter, sometimes used together with cycling, involves seeding aquarium biofilters using biological supplements containing live nitrifying microorganisms, marketed to allow immediate introduction of fish (e.g., Cycle, Hagen Industries; API Quick Start). The effectiveness of these supplements has not been well studied in peer-reviewed investigations, and little is known about supplement microbial community composition and interactions within aquarium biofilters (14). The scientific basis for many of these products comes from literature that predates the discovery of AOA and comammox *Nitrospira* (15–18), which have more recently been shown to dominate freshwater aquarium biofilters (4, 5, 19). Between 9 and 12 million households in the United States maintain a freshwater aquarium representing approximately 140 million individual organisms (20, 21), and the global ornamental home aquarium industry generates an estimated $40 billion CAD per year, most of which represents the freshwater aquarium industry (i.e., fish and associated aquarium equipment) (21–24). The discovery of comammox *Nitrospira* and AOA presents a unique applied microbiology opportunity to develop more effective aquarium supplements with potential for commercial and animal care benefits.

Prior to the discovery of comammox *Nitrospira* and AOA, surveys of aquarium biofilter microbial communities reinforced the importance of *Nitrosomonas* as a central player in nitrogen cycling and ammonia oxidation (15–17, 25). The advent of metagenomic sequencing provided genomic evidence that archaea were also capable of ammonia oxidation (26), which was later confirmed with the isolation of the marine AOA *Nitrosopumilus maritimus* (27). Subsequent aquarium research showed that AOA occasionally dominated AOB in freshwater aquarium biofilters (4, 6). After the discovery of complete ammonia oxidation by comammox *Nitrospira* (28, 29), techniques to detect comammox *Nitrospira* helped demonstrate that they were also abundant in freshwater aquarium biofilters (5, 19, 30, 31). It is now understood that aquarium biofilters rely on a consortium of AOA, AOB, NOB, and comammox *Nitrospira* to maintain low concentrations of ammonia and nitrite, and these consortia establish within the first few weeks of new aquarium operation (19, 32, 33). A recent study demonstrated that comammox *Nitrospira* dominate newly established aquarium filters, regardless of fish load (33). Despite these advances, it is unknown how AOA, AOB, and comammox *Nitrospira* compare when applied as nitrifying supplements.

Formal investigations into using supplements to seed new aquarium biofilters first appeared in literature in the 1970s, when both soil (17) and wastewater effluent (34) were used to promote establishment of a local nitrifying guild. The first biological aquarium supplements containing live nitrifying bacteria were released in the 1980s (e.g., Nutrafin Cycle, Hagen Inc; API Quick Start). Despite their popularity, peer-reviewed investigation into the effectiveness of home aquarium supplements is limited. Since the discovery of AOA and comammox *Nitrospira*, only a single study has investigated the effectiveness of nitrifying supplements intended for home freshwater aquarium systems (14), although this research did not include analysis of the microbial communities and was performed within aquaria lacking fish and organics associated with fish feed and metabolic waste. Nitrifying supplements containing AOA (One and Only, Dr. Tim’s) were included in an assessment of supplements intended for fish transport and associated high ammonia concentrations (35), but were not tested in home aquaria. Contrasted with recent research developing supplements for large-scale high-ammonia recirculating aquaculture systems (36–40), the research gap is apparent. Developing new home aquarium nitrifying supplements containing aquarium-relevant AOA and comammox *Nitrospira* could potentially benefit home aquarium owners, and testing these supplements under aquarium-relevant conditions is needed to compare their effectiveness.

The current work examines the feasibility of using AOA and comammox *Nitrospira* as nitrifying supplements for newly established freshwater aquarium systems, comparing results to an existing nitrifying supplement. Two aquarium-derived enrichment cultures, one dominated by comammox *Nitrospira* (“CX”) and the other enriched with the AOA representative *Ca.* Nitrosotenuis aquarius (“AQ”), were compared against a conventional nitrifying supplement (CS) containing AOB affiliated with the *Nitrosospira* genus. Supplements were applied using a range of doses. Concentrations of ammonia, nitrite, and nitrate were monitored daily, and aquarium filter beads were collected and extracted for genomic DNA to profile the microbial community and quantify nitrifier-specific abundances. Results from this work indicate that newly discovered nitrifiers can serve as supplements for home aquariums given their ability to deplete fish-associated ammonia and nitrite waste while persisting temporally within aquarium biofilters.

## Materials and Methods

### Enrichment cultures and commercial supplements

Dominated by *Ca.* N. aquarius, the supplement AQ enrichment culture was prepared from an aquarium biofilter in the office of Josh Neufeld (University of Waterloo) and has been characterized previously (41). Supplement CX was enriched from material associated with a biofilter sponge (5, 42) from an aquarium biofilter in the office of Josh Neufeld. Both supplements were maintained in 250 mL glass bottles using basal freshwater medium (FWM) containing 0.58 g/L NaCl, 0.15 g/L CaCl_2_•2H_2_O, 0.075 g/L KCl, and 0.05 g/L MgSO_4_•7H_2_O supplemented with 300 µM NH_4_Cl and additional vitamins, buffers, and phenol red as a pH indicator (43). Ammonia concentrations were monitored using Nessler’s reagent (44) and enrichments were amended with an additional 300 µM of NH_4_Cl when ammonia was depleted. Cultures were supplemented periodically with 1 M NaOH to maintain a pH of between 7.0 and 9.0. Both supplements were incubated in the dark without shaking between 28°C and 30°C. The enrichments were propagated into fresh FWM using a 10% inoculum every 12 to 16 weeks. To generate enough volume for experiments, both AQ and CX were scaled to 8 L aerated batch cultures. The supplements were first scaled to 1 L in FWM and maintained for eight weeks before inoculation of 7 L of FWM in 10 L glass bottles equipped with a sterile stainless-steel tube used to bubble air through a 0.22-micron filter. The large-volume batch supplements were maintained in the dark at 30°C, with continuous bubbling, and supplemented with 300 µM NH_4_Cl and NaOH as required. Sterile water was added to offset evaporation (approximately 500 mL per week). The conventional nitrifying supplement (CS) was provided by Hagen Industries.

### Metagenomic sequencing and metabolic characterization of CX

To characterize the genome-encoded metabolic potential of the CX supplement, 2 mL of culture was centrifuged at 10,000×g for 10 minutes and the genomic DNA was extracted from the pellet using a DNeasy PowerSoil Pro Kit (Qiagen). Genomic DNA was submitted to The Centre for Applied Genomics (TCAG) for library preparation and metagenomic sequencing (TruSeq DNA PCR-Free HT Library Prep Kit). The CX sample was sequenced using 20% of a single lane on an S4 flow cell (2x150 cycles) with a NovaSeq 6000 System (Illumina). Raw sequence data were then quality controlled, trimmed, and processed. To generate metagenome-assembled genomes (MAGs), reads were assembled using metaSpades (45), binned using MetaBAT (46), and quality checked and refined using CheckM (47) and DASTool (48), taxonomy assigned using GTDBk (49) and the GTDB r220 database, using the default settings, all contained within the ATLAS v2.19.0 software package (50). Open reading frames were then identified using Prodigal v2.6.3 (51) and annotated by aligning them against the UniRef90 database using DIAMOND v2.1.11 (52).

### *In situ* assessment of nitrifying supplements

To investigate how nitrifying supplements affect aquarium nitrification and nitrifier succession, 33 aquaria were established with 50-watt heaters (Fluval, Hagen Inc.), two plastic plants, an air stone, and an AquaClear 30 side-mounted recirculating filter containing a new sponge, activated carbon filter, and biofilter beads (Fluval, Hagen Inc.) (Figure 1). All materials, excluding biofilter media, were washed with 10% bleach and rinsed with DI water. The aquaria were washed with 10% bleach and rinsed with facility water. The aquaria were then filled with 35 liters of facility water and brought to ∼26°C, after which eight zebrafish (*Danio rerio*) were added to each tank. Of the 33 aquaria, 27 were treated with nitrifying supplements as biological triplicates, 3 remained untreated, and 3 contained no fish. Added supplement was normalized to CS using 16S rRNA gene amplicon sequencing and qPCR amplification copy number based on a simplified treatment scheme of 1× (50 mL), 5× (250 mL), and 10× (500 mL) doses, representing 9.3×10^8^, 4.6×10^9^, and 9.3×10^9^ total nitrifying cells per 35 L, respectively (Table 1). The 1× dose was chosen based on a commercial recommendation for a CS-like supplement. Supplements were introduced to the aquaria above the filter intake immediately after adding fish. Each aquarium was provided with 80 ± 1 mg of Fluval BugBites (Hagen, Inc.) fish food once per day after collecting water samples. Additional facility water was added to each tank weekly to offset evaporation (approximately 2 to 3 liters per week). To test whether nitrifying supplements provide long-term protection against ammonia spikes, the fish were removed after six weeks of operation, and all aquaria were sequentially amended with 10, 30, and 100 µM of ammonium chloride (NH_4_Cl). Aquaria were monitored until ammonia and nitrite were depleted. All experimental protocols were conducted with approval from the Office of Research Ethics at the University of Waterloo (AUPP #46851).

**Figure 1.**
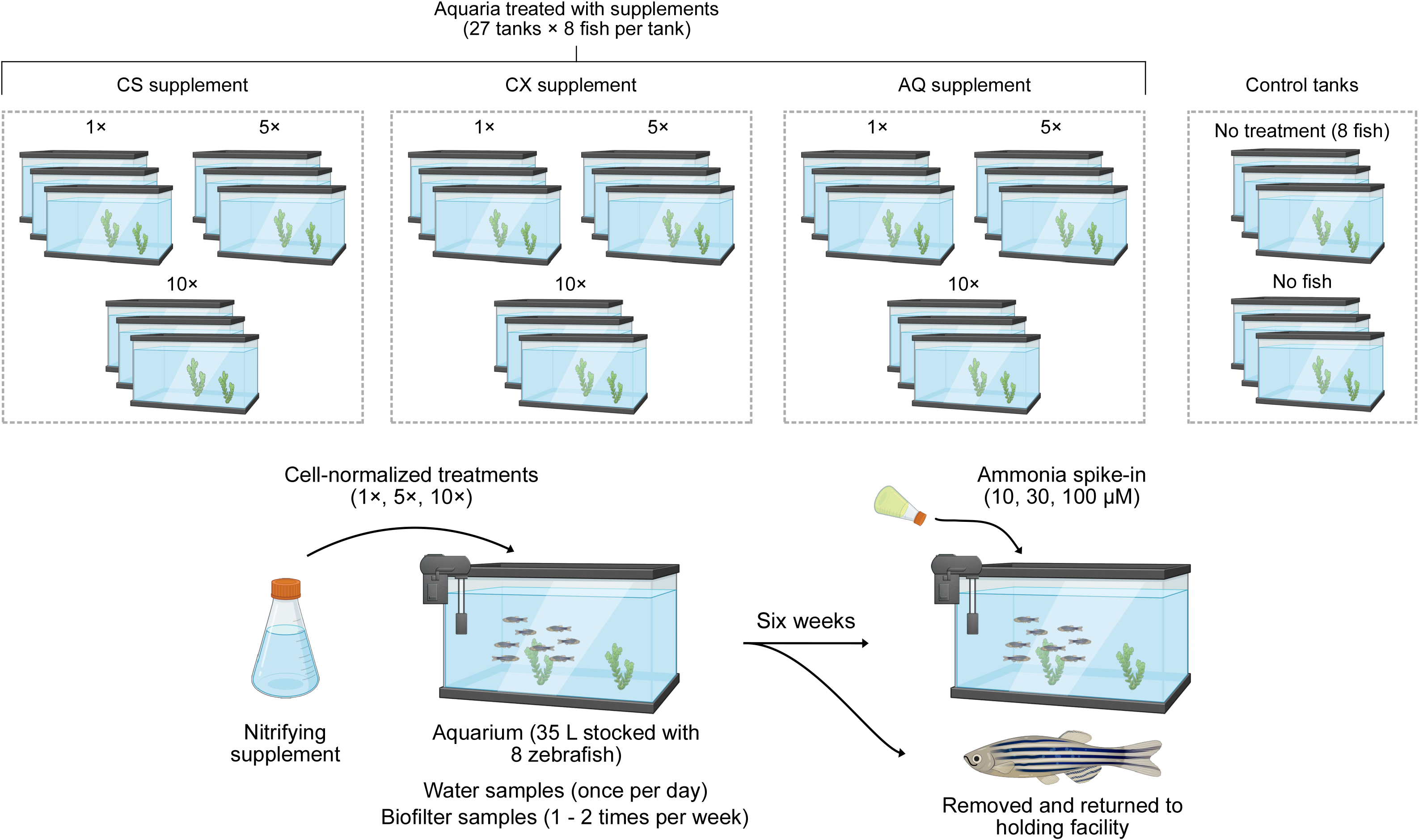
Schematic diagram of the experiment designed to test the *in situ* efficacy of comammox *Nitrospira*, *Ca.* Nitrosotenuis aquarius, and CS as nitrifying supplements.

### Sample collection and processing

Water was collected daily by pipetting directly from the aquarium biofilter and temperature was measured twice a week using a sterilized digital thermometer, in addition to daily monitoring using an adhesive strip aquarium thermometer. Aquarium pH was measured once per week using a general test kit (API Freshwater Master Test Kit), and general water chemistry including chlorine, total hardness, total alkalinity, and carbonate was measured once per week using generic 7-in-1 aquarium test strips. Two biofilter beads were collected twice per week from the center of the biofilter using sterilized tweezers and stored in 10 mL conical tubes at −20°C until processing. At the end of the experiment, a visually determined “clean” and “dirty” 1 cm x 1 cm section of the biofilter sponge was collected for DNA extraction and stored at −20°C until processing. Filtered beads were collected from a shared biofilter from the facility holding tanks (FB), and 1 L of facility water (FW) was passed through a 0.22 µm filter and extracted for genomic DNA. Genomic DNA was extracted from the beads and sponge using a Qiagen PowerSoil DNA extraction kit with a modified protocol to accommodate for the size of biofilter beads, described as follows. The contents of the Power Bead tubes were transferred to a 5 mL conical tube along with 800 µL of the provided “C1” lysis buffer and a single aquarium bead. Tubes were then incubated at 70°C for 10 minutes in a rotating hybridization oven before agitation on a benchtop vortex for 10 minutes at maximum speed. The cell extract was then centrifuged at 7,000 g for 10 minutes, the supernatant transferred to a 1.5 mL tube, and the standard manufacturer protocol resumed.

### Quantification of ammonia, nitrite, and nitrate

Concentrations of total ammonia (NH_3_ and NH_4+_), nitrite (NO_2-_), and nitrate (NO_3-_) were measured for each water sample using OPA (53) and Griess (54, 55) methods in technical triplicate. Water samples were diluted as required for ammonia, nitrite, and nitrate. For ammonia, 100 µL of sample was combined with 100 µL of reagent in a black opaque 96-well plate and incubated in the dark at room temperature for four hours prior to measuring fluorescence at 360/465 nm using a multimode microplate reader (FilterMax F5, Molecular Devices). For nitrite and nitrate, 100 µL of sample was combined with 100 µL of Griess reagent in a clear 96-well plate, and an additional 100 µL of vanadium chloride solution (0.8 g VCl_3_ per 100 mL of 1 M HCl) was added to the nitrate plate. Both plates were then incubated in the dark at 37°C for one hour before measuring absorbance at 550 nm on a spectrophotometer. Serial dilutions for ammonia (50 µM to 0.195 µM), nitrite (100 µM to 0.781 µM), and nitrate (100 µM to 0.781 µM) were used to generate standard curves. Technical triplicates of biological replicates were averaged for each time point.

For each treatment group (n = 10), peak ammonia and nitrite concentrations (n = 30) were tested using one-way ANOVA using base R v4.4.0 and a Tukey post-hoc test using the R package agricolae v1.3-7. For each ammonia spike, zero-order rates for ammonia oxidation were calculated for each aquarium using the slope of the linear portion of the depletion curve (-k), then averaged for each treatment.

### Quantification of AOA, AOB, and comammox *Nitrospira* abundances

To quantify AOA, AOB, and comammox *Nitrospira* populations, *amoA* and 16S rRNA gene copy numbers were quantified in technical duplicate for each bead sample. Samples were diluted to a DNA concentration of between 1 and 10 ng/µL and the *amoA* gene was amplified using the CrenamoA23F/CrenamoA616R (AOA) (56), amoA1F/amoA2R (AOB) (57), and comaAFP/comaARP (comammox *Nitrospira* clade A) (58) primers. Total bacteria and archaea were quantified using the 16S rRNA gene and 515F-Y/806R universal primers (59, 60). Targets were amplified using 15 µL reaction volumes containing 1X SSO SYBR qPCR master mix (7.5 µL of 2X stock) (Biorad), 0.5 µg/µL of BSA, and 2 µL of sample (2 to 20 ng). Depending on the gene target, primers were added at 0.4 µM (AOA or AOB), 0.5 µM (comammox *Nitrospira*), or 0.3 µM (16S rRNA genes). Thermocycler conditions included an initial denaturation at 98°C for 3 minutes, followed by 35 cycles of 98°C for 30 seconds, annealing temperatures of 55°C for 30 seconds (AOA), 60°C for 30 seconds (AOB), 52°C for 45 seconds (comammox *Nitrospira*), or 50°C for 15 seconds (16S rRNA genes), followed by an extension temperature of 72°C for 60 seconds. Sample concentrations were calculated using *amoA*-specific standard curves using representative sequences transformed into plasmids that were subsequently linearized prior to amplification: *Ca.* N. aquarius (AOA, NCBI: KX034182.1), *N. europaea* (AOB, NCBI: L08050.1:249-1079), *Nitrospira inopinata* (comammox *Nitrospira*, NCBI: MN836855.1), and *Thermus thermophilus* for the 16S rRNA gene. Copy numbers were normalized to extraction volume and represent copies per aquarium bead. The limit of quantification for all targets were 100 copies per µL with no obvious signs of off-target amplification. All qPCR curves reported efficiencies of 90 – 110% and *R*^2^ values >0.99.

### Amplicon sequencing and read processing

The V4-V5 region of the 16S rRNA gene was amplified from genomic DNA extracted from filter beads using universal prokaryotic primers 515F-Y (5’- GTGYCAGCMGCCGCGGTAA - 3’) (59) and 926R (5’ - CCGYCAATTYMTTTRAGTTT - 3’) (60) modified to include a 6-base barcode sequence used for identification of amplicons, an adaptor sequence for flow cell binding, and an Illumina primer binding site (61). The PCR amplification was performed as previously described (62) and amplicon sequencing was performed across three separate runs with the 2x250 cycle TG MiSeq Reagent Kit v2 (Illumina Canada, MS-103-1003) using a MiSeq System (Illumina) producing high quality read data (Table S1).

Sequence reads were demultiplexed using the MiSeq Reporter software version 2.5.0.5 (Illumina). Demultiplexed sequences were processed using only the forward reads (single-end) for each amplicon dataset to generate amplicon sequence variants (ASVs) using QIIME2 2024.5 (63) and DADA2 (64). Primers were removed using cutadapt (65) and forward reads were truncated to 224 bases and denoised using DADA2 (64). The ASVs were classified using a naïve Bayes classifier trained against the SILVA 138 SSURef NR99 database (66). Negative controls were manually inspected to ensure that they were distinct from sample profiles. Feature tables and representative sequences from the three sequencing runs were merged using *qiime feature-table merge*, then biological replicates were grouped and summed using *qiime feature-table group*. Finally, feature tables were filtered to remove ASVs with fewer than 10 reads to avoid analysis of spurious sequences generated during sequencing. All sequences generated for the current study were deposited in the European Nucleotide Archive (ENA) under project accession number PRJEB106321.

## Results

### Supplement characterization

Based on 16S rRNA gene profiling, the AQ supplement consisted of *Ca.* N. aquarius as the sole nitrifier, which has been previously characterized (41). The only nitrifier detected in CS was an AOB (i.e., *Nitrosospira*) (Figure 2). The CX supplement was enriched from an aquarium biofilter and originally comprised approximately 30% *Ca.* N. aquarius and 10% *Nitrospira* (circa 2021). The CX supplement has since stabilized and is now enriched with two *Nitrospira* ASVs and AOA are undetectable. Because these *Nitrospira* ASVs represent either comammox or NOB *Nitrospira*, but cannot be distinguished using 16S rRNA gene analysis, the CX supplement was processed for metagenomic sequencing to identify and characterize the *Nitrospira* populations.

**Figure 2.**
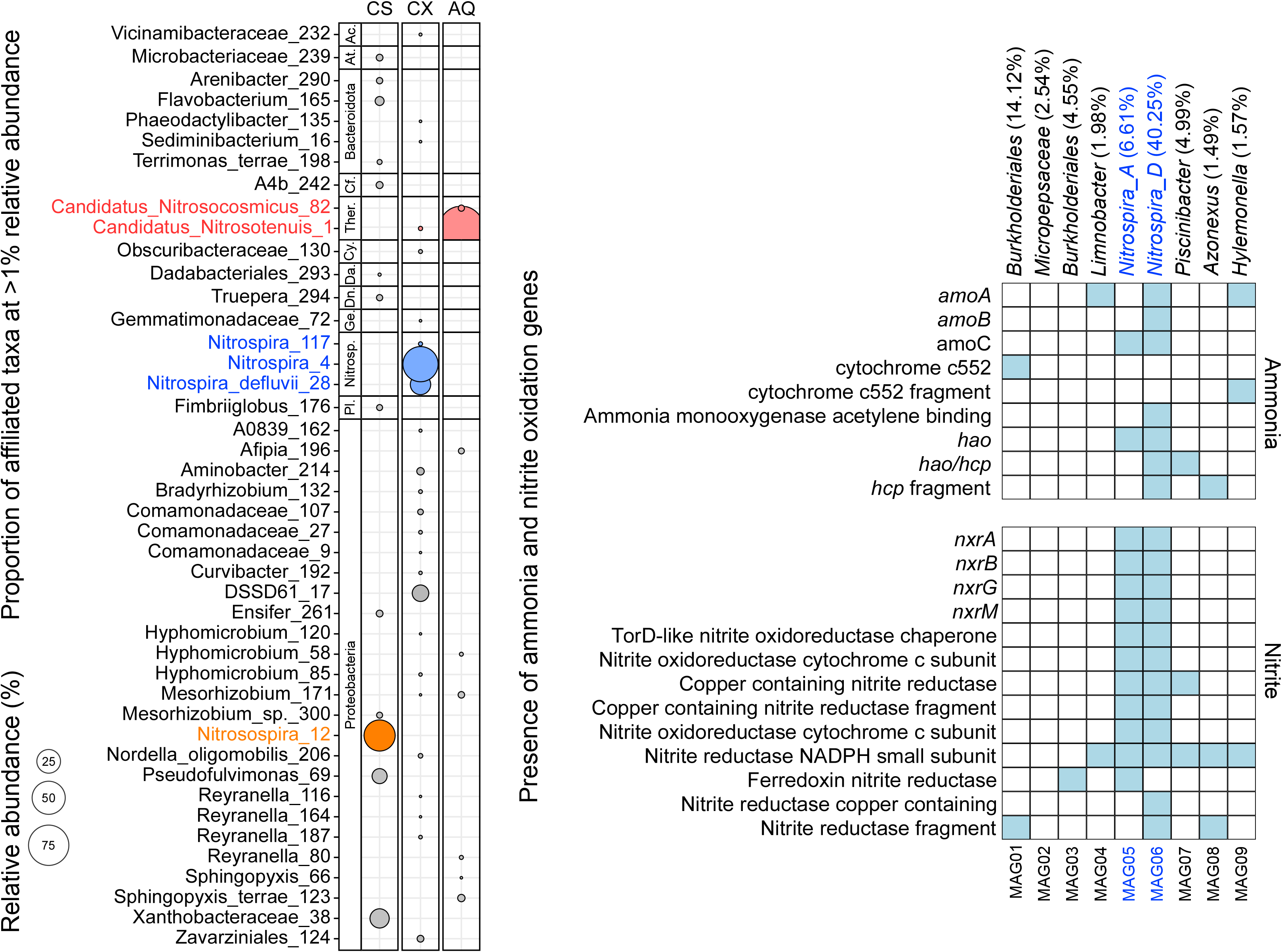
16S rRNA gene profiles of the CS (orange), CX (blue), and AQ (red) supplements (left). Bubble sizes represent relative abundances. Any ASV present at greater than 1% in any sample is shown. Taxa are organized by phylum: *Actinomycetota* (Act.), *Chloroflexota* (Chloro.), *Thermoproteota* (Thermoprot.), *Desulfobacterota* (De.), *Nitrospirota* (Nitrosp.), and *Planctomycetota* (Plancto.). Metabolic characterization of metagenome-assembled genomes constructed from the CX supplement (right). Presence of nitrification-associated genes is indicated by blue squares and is based on alignments of MAG open reading frames to the UniRef90 database. Taxonomic identification and abundance of each MAG in the CX culture are reported above the plot.

Metagenomic analysis of the stock CX supplement produced nine high-quality MAGs (Table S1). Both MAG05 and MAG06 were taxonomically classified as *Nitrospira* yet belonged to distinct lineages based on GTDB classification (Figure 2). All genes required for complete nitrification were detected within MAG06, including subunits for ammonia monooxygenase, hydroxylamine (oxido)reductase, and nitrite oxidoreductases. In contrast, MAG05 lacked ammonia monooxygenase subunits *amoA* and *amoB*, although it contained *amoC* and a hydroxylamine oxidoreductase, as well as nitrite-oxidizing nitrite oxidoreductases and reductases. The remaining MAGs associated with CX did not contain complete pathways associated with either ammonia- or nitrite-oxidation. Based on 16S rRNA relative abundance data and read coverage during MAG construction, MAG06 is likely the dominant *Nitrospira* ASV within the CX supplement and represents a comammox *Nitrospira*, whereas MAG05 likely represents an NOB *Nitrospira*.

### Impact of nitrifying supplements on aquarium nitrification

To test whether treating aquaria with nitrifying supplements improves nitrification rates, new aquaria were treated with 1×, 5×, or 10× doses of CS, CX, or AQ. Increasing the dose reduced ammonia and nitrite accumulation in all aquaria for all three tested supplements (Figure 3). Treating aquaria with 10× doses of CS, CX, or AQ significantly reduced ammonia accumulation compared to untreated aquaria (one-way ANOVA, df = 9, *F* = 8.9, *p* = 2.7×10^-5^). Similarly, nitrite accumulation was significantly reduced for aquaria treated with a 10× dose of CS or CX (one-way ANOVA, df = 9, *F* = 19.2, *p* = 5.3×10^-5^). Aquaria treated with 10× CX had the lowest ammonia and nitrite maxima compared to all other aquaria. Although not significantly different from untreated aquaria, aquaria treated with a 1× dose of CS showed reduced ammonia maxima; nitrite accumulation varied for the 1× and 5× doses. Nitrate accumulated in aquaria treated with CS and the higher nitrate concentrations were associated with aquaria treated with larger doses, which reflected the high concentration of waste nitrate in CS. Treating aquaria with 1× or 5× CX only marginally decreased maximum ammonia concentrations, although nitrite accumulation was significantly reduced for the 5× and 10× dose compared to untreated aquaria. Overall, treatment with any amount of CX maintained nitrite at near or below detectable concentrations. Similar to aquaria treated with CS, nitrate accumulated temporally and differences in total nitrate concentration reflect the concentration of waste nitrate in the enrichment prior to treatment.

**Figure 3.**
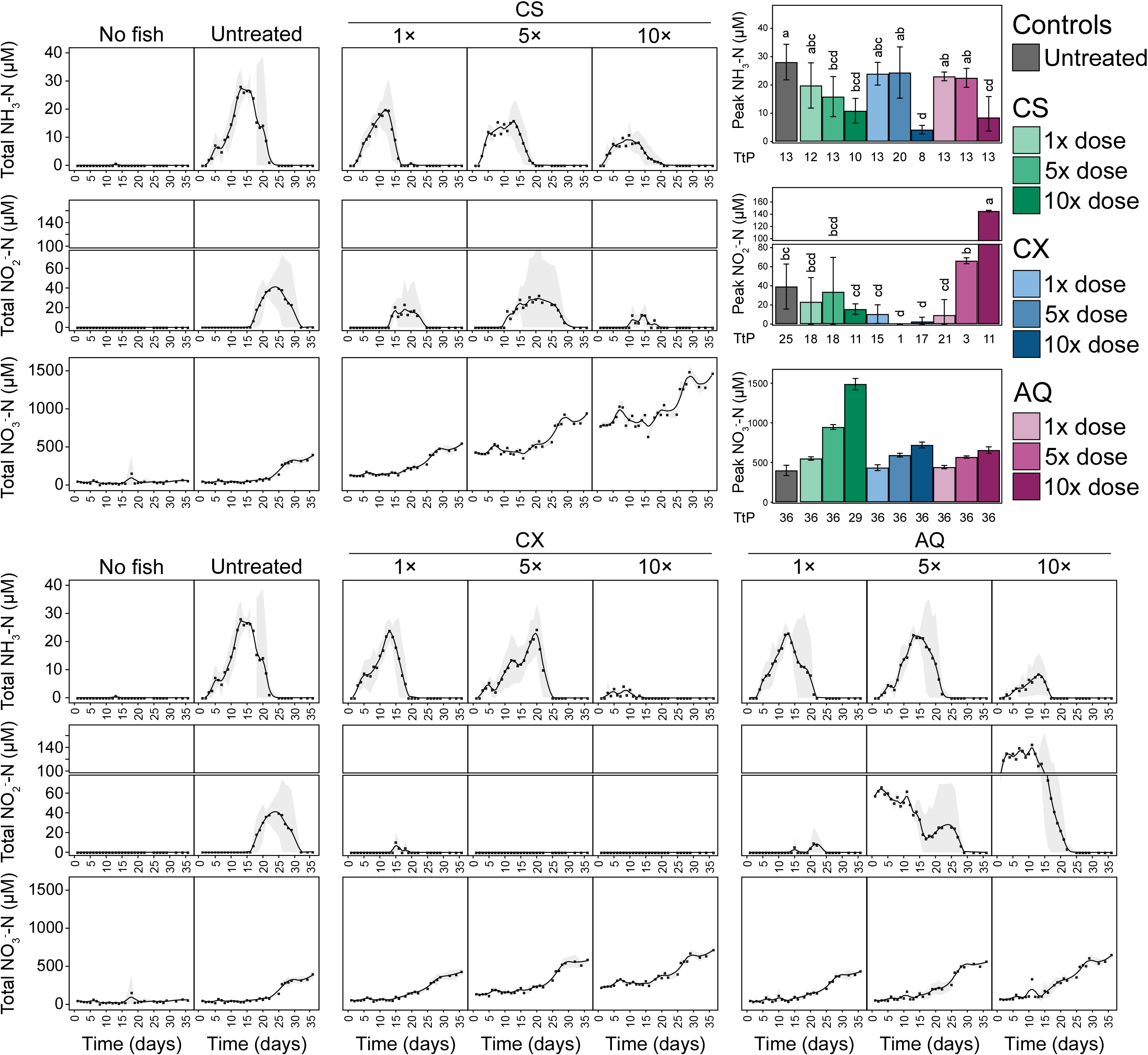
Ammonia, nitrite, and nitrate accumulation in aquaria treated with variable doses of nitrifying supplement. Black squares represent averages of biological replicates (n = 3); grey shading represents standard deviation. A black trendline was calculated using a loess function. Highest detected average concentrations of ammonia, nitrite, and nitrate are shown on the right. Bars represent averages of combined biological and technical replicates and error bars represent standard deviation. Time-to-peak (TtP), representing the sampling time point with the highest measured concentration, is presented along the x-axis of the bar plots and is presented as days. Significant differences in peak concentrations among treatments were tested with a one-way ANOVA and are indicated with letters.

Aquaria treated with AQ were similar to those treated with CX, showing significant reduction in ammonia accumulation when treated with a 10× dose (Figure 3). Treatment with 1× AQ limited the accumulation of nitrite to near detectable limits. Ammonia accumulation for the 5× treatment was similar to the 1× treatment. Waste nitrite present in the AQ stock supplement obscured nitrite accumulation for the 5× and 10× treatments, although the time required for nitrite to be oxidized below detectable limits was comparable to the 1× and 5× CS-treated aquaria, and both were earlier than the untreated control (24 days vs 32 days, respectively).

Aquaria had an average temperature of 26.8°C ± 0.3°C and all aquaria consistently measured between pH 6.5 and 7.1. Aquarium water chlorine (0 mg/L), total hardness (75 – 150 mg/L), alkalinity (120 – 180 mg/L), and carbonate (180 - 300 mg/L) were consistent throughout the experiment for all aquaria.

### Dose dependencies of aquarium filter microbial communities

Microbial community composition was similar for all aquarium biofilter microbial communities (Figure 4). Sequences affiliated with *Limnobacter* and *Aquabacterium* were abundant in the first week of aquarium operation, followed later by *Comamonadaceae*, *Ferrovibrio*, *Methyloversatilis*, and *Amphilicatus*, among others. Several ASVs associated with *Planctomycetota* were detected in all aquaria in the second week but absent in the final weeks. ASVs associated with *Nitrospira* were consistently detected after approximately two weeks and persisted at high relative abundance for the remainder of the experiment. An ASV associated with *Ca.* Nitrosotenuis in the AQ-treated aquaria was detected on day 0 and increased to represent a relatively large proportion of the total microbial community (0.01% to 67.1%). Microbial communities within biofilter sponges differed from those on the beads, represented by

**Figure 4.**
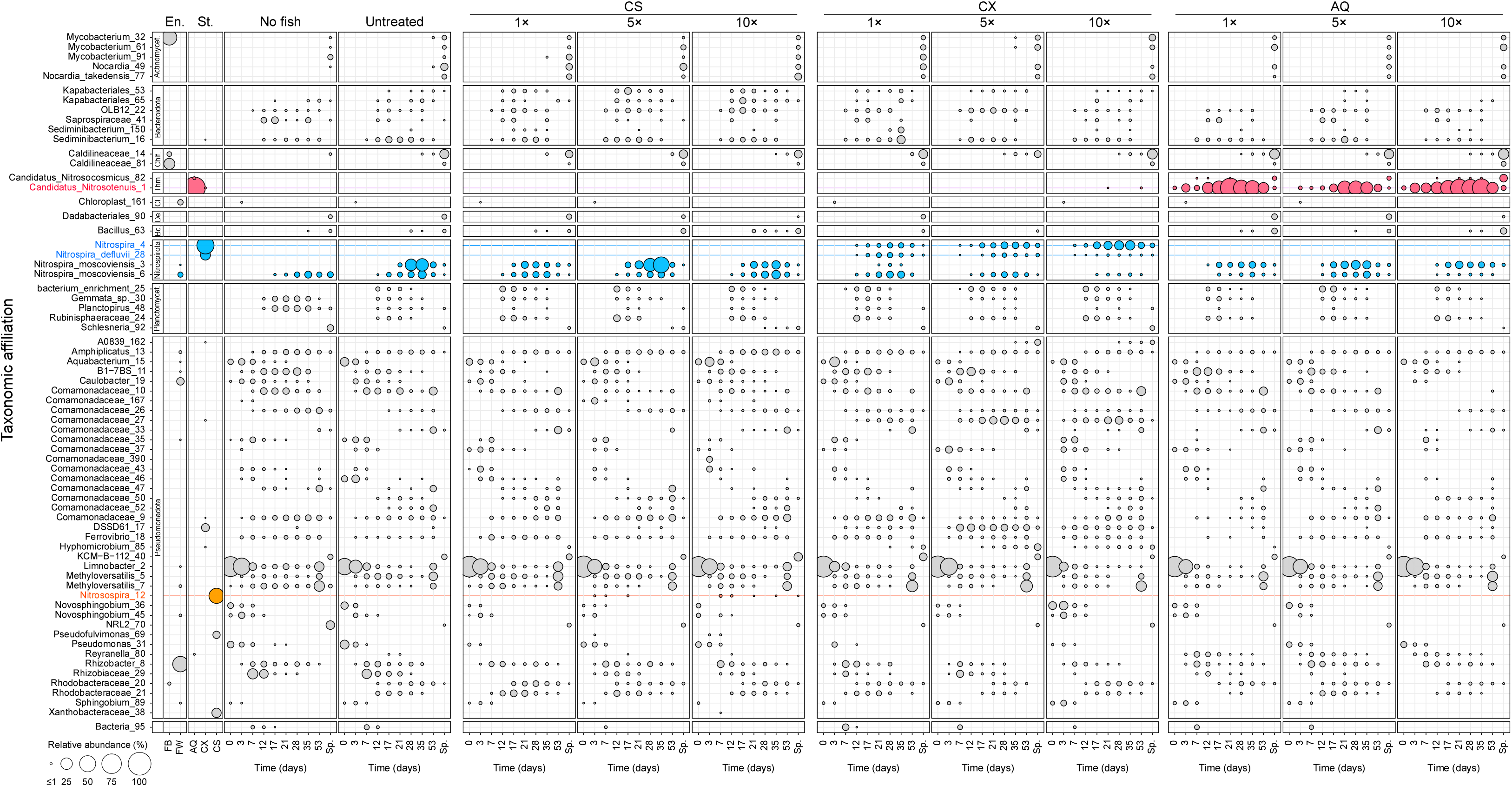
Relative abundance and distribution of 16S rRNA ASVs within aquarium biofilters treated with nitrifying supplements. ASVs associated with CS (orange), CX (blue), and AQ (red) are indicated with a coloured line. Bubble area represents the relative abundance of an ASV within a sample, and only ASVs present at >5% in any sample are shown. Environmental references include “FB” (fish housing filter beads) and “FW” (facility water). Taxa are organized by phylum categories: *Actinomycetota* (Actinomycet.), *Bacteroidota, Chloroflexota* (Chlf.), *Thermoproteota* (Thm.), Chloroplast (Cl.), *Desulfobacterota* (De.), *Bacillus* (Bc.), *Nitrospirota, Planctomycetota* (Planctomycet.), *Pseudomonadota*, and unclassified *Bacteria*.

ASVs affiliated with *Mycobacterium*, *Caldilineaceae*, and an ambiguously classified *Acidithiobacillaceae* (ASV KCM-B-112_40). Aquaria stocked with fish contained microbial taxa distinct from aquaria without, including *Nocardia* and *Rhodobacteraceae*, as well as several ASVs associated with *Comamonadaceae*. Few relatively abundant taxa (e.g., >5% in any sample) were uniquely associated with profiles from specific treatments. Biofilters treated with CS contained unique ASVs affiliated with *Pseudofulvimonas*, whereas CX-treated aquaria contained an exclusive *Rhizobiales*-affiliated ASV, A0839_162.

Nitrifier populations detected in aquarium biofilter profiles contained representatives common to all aquaria, as well as distinct ASVs unique to each treatment (Figure 5). In all aquaria, except for those treated with AQ, ASVs affiliated with *Nitrospira* were dominant among putative nitrifiers, with total relative abundances exceeding 25% of the total microbial community for several samples. *Nitrospira*-associated ASVs were prevalent within all sample profiles, including within biofilter beads collected from aquaria where the fish were held prior to the start of the experiment (i.e., sample “FB”). A *Nitrospira*-associated ASV (Nitrospira_moscoviensis_6) detected in the source water used to fill the aquaria (i.e., sample “FW”) persisted in all aquarium biofilter profiles. A second *Nitrospira* ASV (Nitrospira_moscoviensis_3) associated with source water was also prevalent in many biofilters at relatively high abundance, although mostly absent in aquaria treated with CX. Aquaria treated with CX contained *Nitrospira* ASVs unique to the CX supplement. Specifically, the ASV Nitrospira_4 increased in relative abundance with higher CX doses, whereas Nitrospira_defluvii_28 decreased. A *Nitrospira*-associated ASV, Nitrospira_117, was detected in all fish-containing aquaria and associated with the facility holding tanks. Aquaria treated with AQ were dominated by *Ca.* N. aquarius, in several samples reaching relative abundances of greater than 60% of the total microbial community and increasing in relative abundance with higher doses. Absolute abundances of AOA were also highest for samples collected from aquaria treated with AQ. The AOB were at low relative abundance for all aquaria, although aquaria treated with CS contained the highest absolute AOB abundances. The CS-associated nitrifier, Nitrosospira_12, was detected for several samples associated with two of the three CS dose groups (i.e., 5× and 10× CS treatment) and was unique to aquaria treated with CS. Otherwise, AOB were represented by a Nitrosomonas_ureae_103 ASV, as well as DSSD61_17 associated with *Nitrosomonadaceae*, and prevalent in aquaria treated with CX.

**Figure 5.**
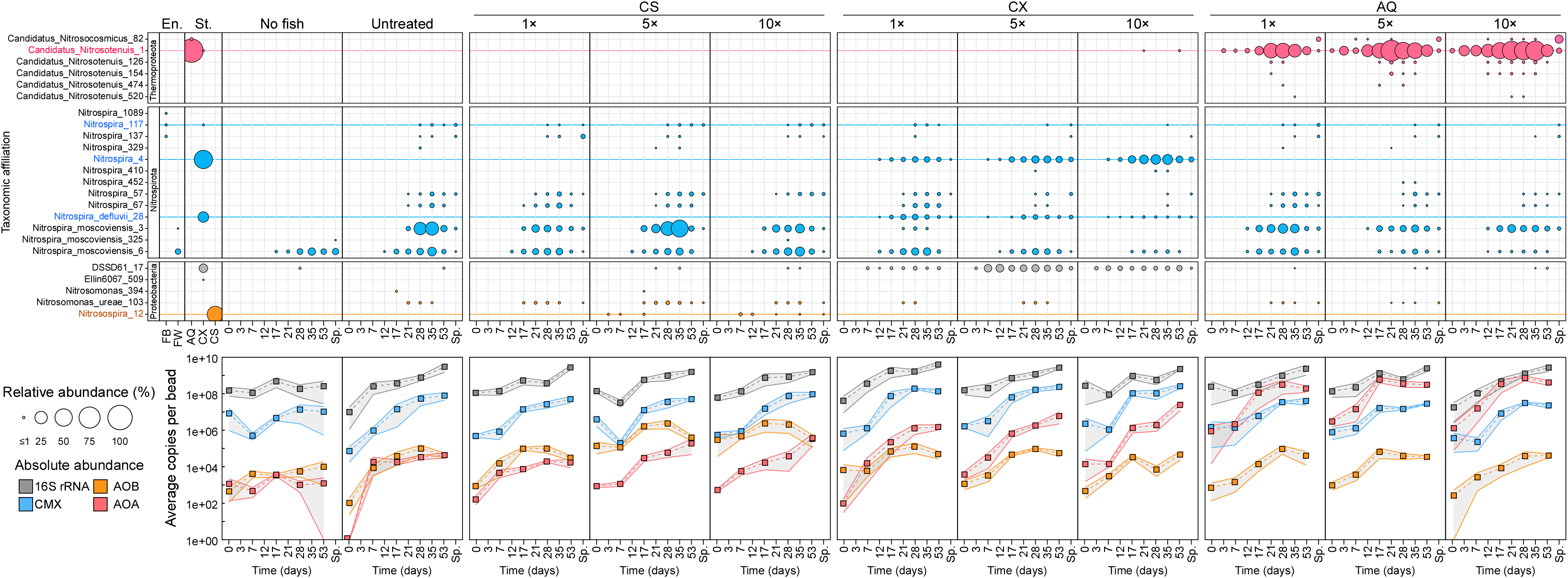
Nitrifier relative and absolute abundance within aquaria treated with variable supplement doses. ASVs associated with CS (orange), CX (blue), and AQ (red) are indicated with a coloured line. Bubble area represents the relative abundance of an ASV within a sample, and only ASVs present at >0.1% in any sample are shown (top panel). Environmental references include “FB” (fish housing filter beads) and “FW” (facility water). Absolute abundances based on nitrifier-specific *amoA* gene qPCR amplification (bottom panel). Squares represent averages of biological replicates (n = 3) and grey shading represents standard error of the mean.

### Ammonia spikes

All fish were removed after six weeks and aquaria were amended with 10, 30, and 100 µM of NH_4_Cl “spikes”. For the 100 µM spike, the mean ammonia oxidation rate in aquaria treated with a 10× dose of CS was significantly higher compared to aquaria treated with 10× CX or AQ but was comparable to aquaria treated with 5× CX or AQ (one-way ANOVA, df = 9, *F* = 26.5, *p* = 2.0×10^-16^) (Figure 6, supplemental material). Despite detected differences among treatments, the magnitudes of the differences were small. Among all spike concentrations, aquaria showed variations in ammonia oxidation rates seemingly independent of the applied nitrifying supplements.

**Figure 6.**
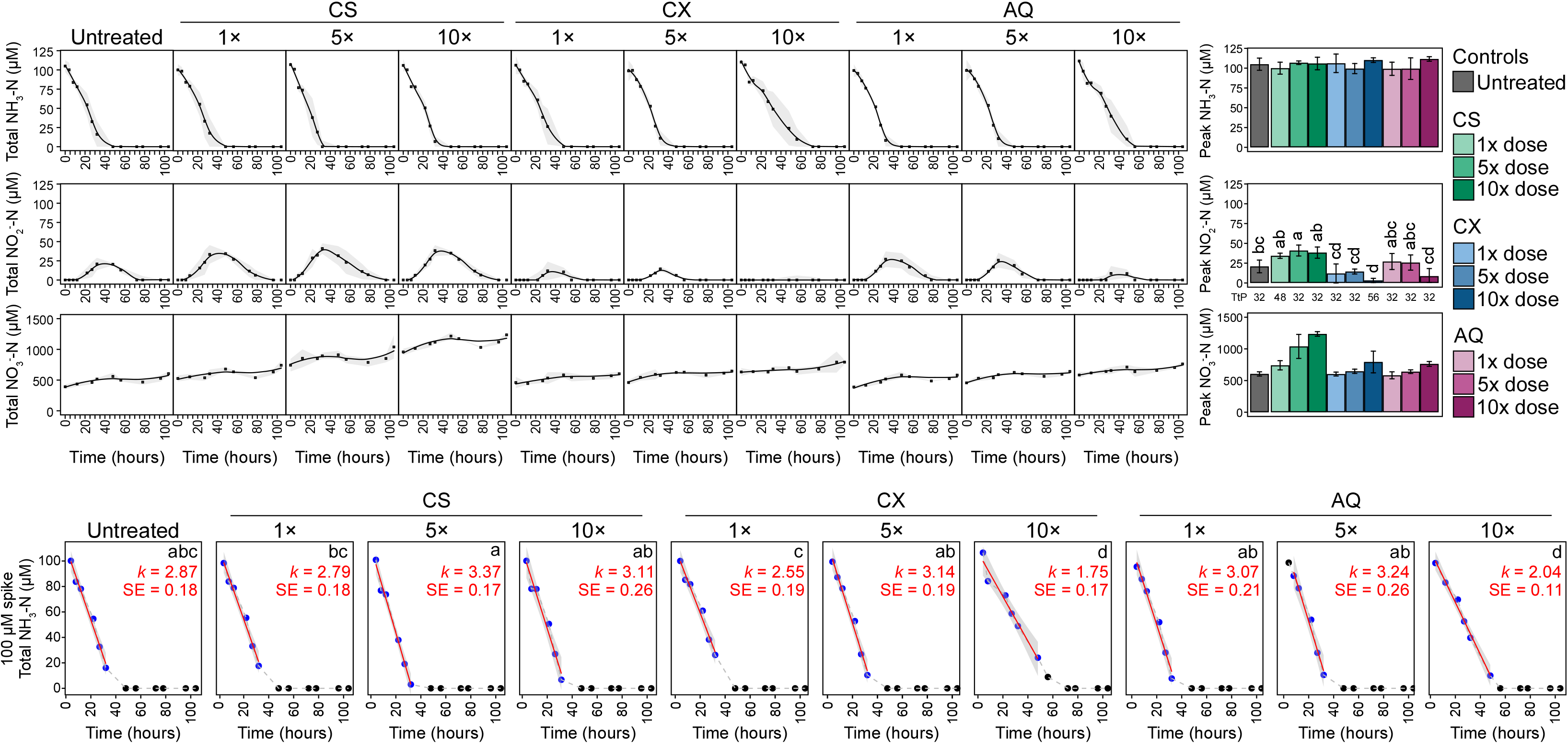
Ammonia, nitrite, and nitrate concentrations following additions of 100 µM (top). Black squares represent averages of biological replicates (n = 3); grey shading represents standard deviation. The average highest detected concentrations for each treatment are shown to the right with error bars representing standard deviation. Time-to-peak (TtP) for nitrite, representing the sampling time with the highest measured concentration, is presented along the x-axis of the bar plots and is represented in hours. Significant differences in peak concentrations among treatments were tested with a one-way ANOVA and are indicated by letters. Total aquarium ammonia oxidation rates following spiking with 100 µM of ammonium chloride (bottom). Blue circles indicate the data used to calculate zero-order rates. A red line indicates the mean -slope of the curve with mean rate calculated as *k* (µM/hr) (n = 3). Significant differences (*p* value < 0.05) calculated with a one-way ANOVA are indicated with letters.

Nitrite accumulation was significantly reduced in aquaria treated with 10× CX for the 100 µM treatment, whereas nitrite maxima were significantly higher in aquaria treated with 5× CS (one-way ANOVA, df = 9, *F* = 11.8, *p* = 3.2×10^-6^) (Figure 6). Aquaria treated with any dose of CX consistently had the lowest peak nitrite concentrations across all ammonia concentrations. Aquaria treated with CS often had the highest peak nitrite concentrations, whereas those treated with AQ showed variable results.

## Discussion

### Dose dependency of nitrifying supplements

Applying nitrifying supplements to newly established biofilters is intended to improve early nitrification rates, facilitate safe introduction of fish into new aquaria, and protect against ammonia and nitrite toxicity (9, 10). Nitrifying activity was improved by supplement addition for all treatment conditions, with reductions in ammonia and nitrite accumulation most apparent for aquaria treated with a 10× dose (Figure 3; see Materials and Methods for a description of doses). Treating aquaria with CX maintained nitrite near or below detectable limits for all doses, and ammonia concentrations remained near detectable limits in aquaria treated with a 10× dose. The difference in early ammonia oxidation activity between the 5× and 10× dose may reflect a cell abundance threshold of between 4.6×10^9^ and 9.3×10^9^ nitrifying cells per 35 L, which enabled ammonia to be oxidized faster than it was produced under the current tested fish load.

Profiling of nitrifiers within CX-treated aquarium biofilters showed the relative abundance of the ASV Nitrospira_4 increased with dose, whereas other *Nitrospira* ASVs decreased in relative abundance. These additional *Nitrospira* ASVs were associated with untreated aquarium biofilters and facility water, which were likely displaced by more competitive *Nitrospira* populations within CX, particularly in aquaria treated with higher doses of CX where adding increased cell numbers conferred an additional competitive advantage. The ASV Nitrospira_4 represents the *Nitrospira* population that persists at approximately 60% relative abundance in the CX supplement and is presumably associated with comammox activity (Figure 2). Thus, the actual ammonia-oxidizing-cell threshold to eliminate ammonia accumulation may be lower. Additionally, Nitrospira_4 was detected only in CX-treated biofilters, suggesting that improved nitrifying activity observed in CX-treated aquaria resulted from the introduction of *Nitrospira* members unique to the CX supplement. The CX supplement contains a consortium of NOB and comammox *Nitrospira* that can completely oxidize ammonia to nitrate. Although contributions of the respective *Nitrospira* populations were not quantified in this study, both NOB and comammox *Nitrospira* likely contributed to reduced nitrite maxima by either oxidizing nitrite produced by existing ammonia oxidizers or through bypassing nitrite release entirely using comammox activity. Regardless, the CX supplement is a consortium that improves ammonia and nitrite oxidation and mitigates accumulation, seeming to perform best when applied to new aquaria at concentrations of at least 9.3×10^9^ total nitrifying cells per 35 L (∼2.6×10^8^ cells per L), which was delivered by adding ∼1 L of CX supplement into each aquarium.

Only a 10× dose of AQ was sufficient to observe significant differences in ammonia accumulation compared to untreated aquaria (Figure 3). However, both relative and absolute AOA abundances were higher compared to comammox *Nitrospira* in AQ-treated aquaria. Considering that *Ca.* Nitrosotenuis was the sole nitrifier detected within the AQ supplement, presumably representing all of the added 9.3×10^9^ nitrifying cells, these results may suggest that *Ca*. Nitrosotenuis has a lower per-cell activity within this system compared to the CX-associated comammox *Nitrospira*. Although nitrite present in the AQ supplement masked nitrite accumulation in AQ-treated aquaria, increases in nitrite concentrations between the 15- and 25-day sampling timepoints suggest accumulation. Data for the 1× and 10× AQ treatments suggest that nitrite oxidation was improved, despite AQ containing no nitrite oxidizers.

The natural succession of a local nitrifying guild within new aquaria would typically first require ammonia oxidizers, followed by nitrite oxidizers when nitrite becomes available. However, the AQ supplement contained waste nitrite, which when added to the aquaria likely enriched existing NOB populations present in the facility water and associated with the fish. Nitrite is less toxic to fish compared to ammonia but may still result in illness or death with extended exposure (67–69), requiring careful attention during new aquarium startup. Nitrite concentrations were not increased to toxic concentrations by the addition of AQ and no fish deaths or changes in fish behaviour were observed throughout experiment. Instead, AQ reduced ammonia accumulation while also priming for NOB activity, suggesting that an AOA-based aquarium supplement containing *Ca.* N. aquarius could be effective for new aquarium application.

Previous studies have shown variable presence (4, 19) or absence (33) of archaea in aquarium biofilters, suggesting they may be lacking from environmental reservoirs and require seeding. The relative absence of detectable AOA in the facility water, beads, and non-AQ aquarium biofilters demonstrates that environmental sources for this experiment were insufficient for inoculating biofilters with AOA. However, these experiments demonstrate that AOA persist in biofilters when intentionally introduced using nitrifying supplements and additionally appear capable of displacing comammox *Nitrospira* as dominant ammonia oxidizers in freshwater aquaria (Figure 5).

Aquaria treated with CS demonstrated incremental improvements in ammonia oxidation activity with increased dose (Figure 3). Peak average ammonia concentrations decreased with increased dose. The *Nitrosospira* ASV associated with CS was detected for several samples but only at low relative abundances, and only for aquaria treated with 5× and 10× CS. Poor establishment of the CS-associated *Nitrosospira* may have resulted from weak competition against existing nitrifying populations in relatively low-ammonia conditions. Additionally, selective pressures favouring high affinities for ammonia would be present following depletion of any accumulated ammonia, potentially excluding the low-affinity AOB (43) and hindering additional establishment. Indeed, absolute abundances of AOB increased with increased dose, then generally decreased in the following days and weeks post-accumulation. The increase in absolute abundances of AOB did not coincide well with relative abundances of the *Nitrosospira* ASV dominant in CS. Instead, an ASV associated with *Nitrosomonas ureae* was detected among all CS-treated aquaria, as well as several other aquaria, but was not associated with the stock CS profile. Thus, the increased ammonia oxidation activity observed for higher doses of CS may represent temporary activity associated with the population of *Nitrosospira* that did not establish well within the biofilters but may have persisted in circulating aquarium water at least temporarily.

Compared to the CX supplement, CS provided less consistent protection against nitrite accumulation. Aquaria required a 10× dose of CS to achieve nitrite oxidation effects similar to the 1× CX dose. Additionally, both the 1× and 5× CS doses showed high variability among biological replicates, whereas the 10× dose showed more consistent nitrite oxidation. Although these results imply that treating aquaria with more CS can improve nitrite oxidation activity, the lack of detectable NOB within the CS stock and variability among replicates and treatments suggest that environmental factors and reservoirs may have a relatively strong influence. Nitrifiers are known inhabitants of premise plumbing and water treatment systems (70–73) and several *Nitrospira* ASVs were detected in water used to fill the aquaria (Figure 5). Additionally, fish were housed within a holding system with a shared biofilter prior to the experiment, following a separate experiment investigating CX activity. Based on 16S rRNA gene profiles, the facility holding tanks contained a *Nitrospira* ASV associated with CX (Nitrospira_117). Thus, resident populations of *Nitrospira* in plumbing and holding tanks acted as reservoirs by which all aquaria were inoculated, likely contributing to the overall abundance of *Nitrospira*. Nitrospira_117 was detected among all aquaria and, in combination with other non-CX associated *Nitrospira* present in the facility, may be responsible for some of the nitrite oxidation observed in CS-treated aquaria.

### Ammonia treatments

Ammonia and nitrite accumulation is not just a concern for the first several weeks but for the entire life of an aquarium, which may be in operation for many years. Acute ammonia loading events caused by excessive feeding, sudden degradation of organic matter (e.g., dead fish), or treatment of sick fish (e.g., antibiotics), can increase ammonia and subsequent nitrite concentrations. Nitrifying supplements that provide both immediate and long-term improvements to nitrification activity can provide aquarium owners with an additional tool to protect against toxicity-related fish stress.

Overall, adding nitrifying supplements to aquaria during initial startup produced only small differences in ammonia oxidation rates following an acute ammonia loading event (Figure 6, supplemental material). Although adding higher doses of supplement may provide sufficiently abundant populations of nitrifiers to prevent initial ammonia accumulation, local nitrifier guild composition and abundance appear to be defined by initial aquarium ammonia loading. Considering that all aquaria were under identical ammonia and nutrient loads, the observed ammonia oxidation rates likely reflect the maximum activities capable of being supported within this current system. Potential differences in ammonia oxidation rates among supplement-specific nitrifiers, such as comammox *Nitrospira* and *Ca.* N. aquarius, may be equalized among aquaria via changes in population size due to resource limitations shared across all aquaria, producing similar total aquarium ammonia oxidation rates.

Whereas similar ammonia oxidation rates among aquaria may show the effect of resource limitations and maximum carrying capacity, differences in nitrite accumulation following acute ammonia loading highlight specific impacts of supplement treatments. Treatment of aquaria with CX established nitrifying populations that protected against nitrite accumulation (Figure 3, Figure 6). Increasing the initial dose of CX improved this protective effect, preventing nitrite accumulation under all spike concentrations. The long-term protective benefits of CX appear to be linked to two *Nitrospira* ASVs unique to the CX culture and only observed within aquaria treated with CX: Nitrospira_4 and Nitrospira_defluvii_28. Metagenomic analyses of the CX supplement suggest that Nitrospira_4 represents a comammox *Nitrospira* (Figure 2) and, given that it was the most abundant *Nitrospira* ASV in CX-treated aquaria, the long-term protection provided by CX appears mostly due to comammox activity that did not result in any nitrite accumulation. This is reinforced by the observation that Nitrospira_4 was at higher relative abundances in aquaria treated with 10× CX where no nitrite accumulation was observed following acute ammonia loading. Additionally, the NOB *Nitrospira* associated with the CX supplement may also contribute to the elimination of nitrite produced from ammonia oxidizers. Thus, treating aquaria with CX during initial aquarium startup populates aquarium biofilters with *Nitrospira* populations that confer better protection against sudden nitrite accumulation following acute ammonia loading events.

With higher doses of CS, nitrite accumulation was greater compared to the untreated aquaria for all ammonia spike concentrations (Figure 6). It is unclear why the untreated aquaria might accumulate less nitrite following ammonia spiking because both the untreated and CS-treated aquaria contained relatively similar *Nitrospira* profiles. The CX-derived Nitrospira_117 was unintentionally introduced into facility holding tanks prior to the experiment and thus acted as an environmental reservoir of potential comammox *Nitrospira*. This ASV was present across all aquaria and absolute abundances of comammox *Nitrospira* were slightly higher in the untreated aquaria compared to CS (Figure 5), suggesting that a larger proportion of ammonia may have been oxidized directly to nitrate. Introducing a population of AOB could have influenced the succession of NOB populations by producing more nitrite earlier compared to the untreated aquaria. Additionally, any undetected NOB within CS may have occupied niches in the biofilter and prevented establishment of resident *Nitrospira*, which perhaps have characteristics enabling slightly faster nitrite oxidation rates.

As might be expected, treating aquaria with AQ resulted in more nitrite accumulation after spiking with ammonia (Figure 6). Unexpectedly, increasing the dose of AQ produced lower nitrite concentration maxima following spiking with ammonia, and nitrite remained near detectable limits in aquaria treated with a 10× dose of AQ. In this case, comammox *Nitrospira* populations were lower based on qPCR amplification absolute abundance and there are no clear explanations for differences in nitrite oxidation based on 16S rRNA gene profiles. One major difference in the AQ-treated aquaria is the introduction of nitrite during initial inoculation, nearing 140 µM of nitrite for the 10× dose (Figure 3). This may have enriched existing NOB populations, and the increased population size enabling faster nitrite oxidation rates may not be observable within the relative abundance data.

## Conclusion

New aquarium biofilters lack a sufficiently abundant local nitrifying guild capable of oxidizing ammonia and nitrite, and the subsequent accumulation of these compounds may become toxic to housed fish. Nitrifying supplement cultures of aquarium-relevant *Ca.* N. aquarius or comammox *Nitrospira* were effective at establishing within aquarium biofilters and improving nitrification. The results of this study show that adding nitrifying supplements to new aquaria prevented ammonia and nitrite accumulation when added at high doses associated with a nitrifying cell abundance of between 4.6×10^9^ and 9.3×10^9^ cells per 35 L. Treating new aquaria with 10× CS, CX, or AQ significantly reduced the peak ammonia concentrations compared to untreated aquaria, and treating with 10× CX consistently prevented ammonia and nitrite accumulation. Treating aquaria with CX also provided protection against nitrite accumulation following acute ammonia loading events. Together, these results suggest that CX would be effective as a nitrifying supplement for use in freshwater aquaria.

Microbial community profiling and targeted *amoA* qPCR amplification showed that biofilters treated with CS, CX, or AQ contained nitrifying populations unique to each treatment. Comammox *Nitrospira* were dominant in biofilters treated with CS and CX, whereas AOA were dominant in biofilters treated with AQ, and AOB abundance was relatively low in all aquarium biofilters. The distribution of nitrifying populations among aquarium biofilters demonstrates the deterministic nature of aquarium biofilter microbial community development and the ecological adaptations of specific nitrifying populations to low-ammonia environments. Additionally, these results show that environmental reservoirs alone may be insufficient for introducing specific nitrifying populations, such as the AOA, into aquarium biofilters. Intentionally introducing aquarium-relevant nitrifiers to new aquaria results in their persistence within biofilters, and inoculating new aquaria with nitrifying supplements is recommended to promote the development of effective nitrifying populations that protect fish against ammonia and nitrite toxicity.

## Acknowledgments

We thank Wayne J. Parker for helpful suggestions related to interpretation of rate data and Barbara J. Butler for discussions and advice. Additionally, thank you to Nicholas Bendo and Martin Ryan at the University of Waterloo’s WATER facility for animal care support, and to Rachel Beaver for sequencing services. This work was funded by an Alliance grant from the Natural Sciences and Engineering Research Council of Canada (NSERC) in partnership with Hagen Industries (Saint-Laurent, Quebec, Canada).

## Table and figure legends

Table 1. Normalization of supplement treatment volumes based on total nitrifying cell abundances. Both CX and AQ were normalized to CS using a 50 mL dose as recommended by the CS supplement. Nitrifying cells were estimated using qPCR targeting the 16S rRNA gene combined with 16S rRNA relative abundance data and proportions of known nitrifiers.

